# Connecting the dots between computational tools to analyse soil-root water relations

**DOI:** 10.1101/312918

**Authors:** Sixtine Passot, Valentin Couvreur, Félicien Meunier, Xavier Draye, Mathieu Javaux, Daniel Leitner, Loïc Pagès, Andrea Schnepf, Jan Vanderborght, Guillaume Lobet

## Abstract

In the recent years, many computational tools, such as image analysis, data management, process-based simulation and upscaling tools, were developed to help quantify and understand water flow in the soil-root system, at multiple scales (tissue, organ, plant and population). Several of these tools work together or, at least, are compatible. However, for the un-informed researcher, they might seem disconnected, forming a unclear and disorganised succession of tools.

In this article, we present how different pieces of work can be further developed by connecting them to analyse soil-root-water relations in a comprehensive and structured network. This “explicit network of soil-root computational tools” informs the reader about existing tools and help them understand how their data (past and future) might fit within the network. We also demonstrate the novel possibilities of scale-consistent parameterizations made possible by the network with a set of case studies from the literature. Finally, we discuss existing gaps in the network and how we can move forward to fill them.

**Highlights:** Many computational tools exist to quantify water flow in the soil-root system. These tools can be arranged in a comprehensive network that can be leveraged to better interpret experimental data.

## Water flow in the soil-root system

Water deficit is one of the most dramatic abiotic stresses in agriculture [8]. It occurs when leaf water supply is limited by either the low potential of soil water, and/or by the high hydraulic resistance of the soil-plant system [9]. At this point, the atmospheric demand for water is hardly met and stomata close, reducing the plant transpiration and photosynthesis. To investigate when such limitation occurs, the complex plant-soil-atmosphere system is often conceptualised as a multidimensional hydraulic network, in which both soil and root hydraulic properties may substantially control shoot water supply [10–12].

The **structural properties** of the roots compose the first dimension of the soil-root hydraulic network. Structural properties refer to the physical position and arrangement of the objects of interest. They can be conceptualised at the tissue/organ (transversal anatomy, fig. 1A), plant (root architecture, fig. 1B), or population scale (rooting density profile, fig. 1C).

**Figure 1:**
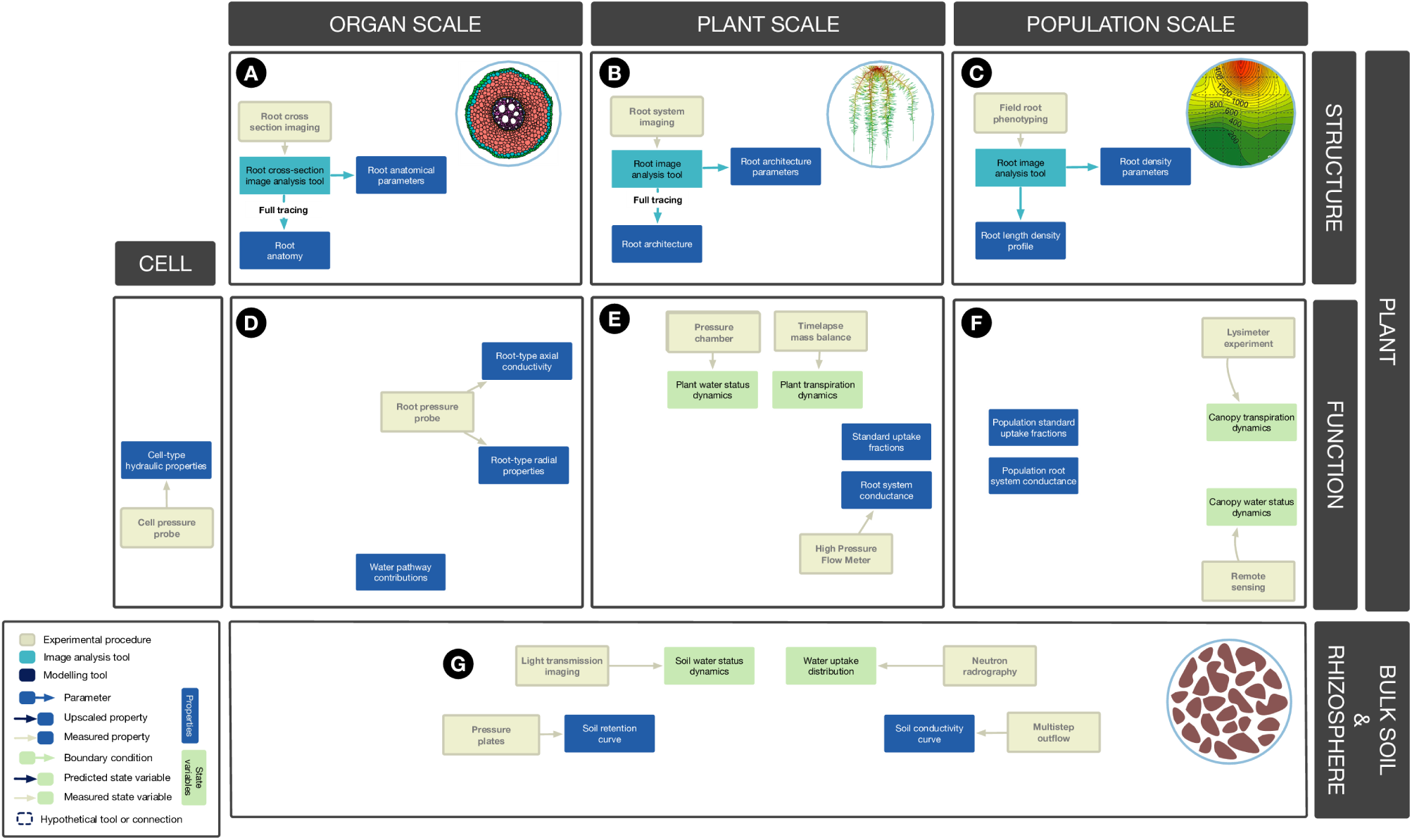
Quantifying water relations in the soil plant system. Tools, properties and state variables used to quantify: **(A)** the structure of root organ; **(B)** the structure of root system; **(C)** the structure of root profiles; **(D)** the water flow at the organ scale (root section); **(E)** the water flow at the plant scale (root system); **(F)** the water flow at the population scale; **(G)** the water flow in the soil. Without appropriate tools, variables of interest, scales and even plants and their environment seem disconnected.

A second dimension, overlaying structural properties, encompasses the system **functional properties**. When studying water movement, functional properties often refer to hydraulic conductivities or reflection coefficients. Like in the structural layer, these properties can be defined at different scales. Local radial and axial hydraulic conductivities can be defined at the organ scale (fig. 1D) while the entire root system of a single plant can be characterized by its conductance (fig. 1E) and would relate to plant water status [13]. An extension of this property to the population scale is the plant population hydraulic conductance per unit horizontal area (fig. 1F), common in canopy models [14], and recently integrated in root models [15].

Finally, a third dimension describes the plant **environment**. In this contribution, we focus on the soil compartment, which includes the rhizosphere and the bulk soil (fig. 1G). Their respective spatial domains are concentric around individual roots, and their properties differ substantially, so that the rhizosphere is often considered to critically affect plant water availability under water deficit [16,17]. The bulk soil and rhizosphere hydraulic properties may be described by their water retention and hydraulic conductivity curves. The former defines the pressure needed to extract water from the porous media, and the latter the relation between water flux and water potential gradient in space [18]. The water potential that defines the energy level of water is a critical environmental variable, driving the flow of water in the soil-plant system. Similarly to the plant, the soil could be divided into functional and structural components and described according to the studied scale. However, we did not explicit this separation in the following as we rather focus on the plant property description in this study.

Each element of the network is dynamic and heterogeneous. Root systems grow, develop and take up water, while soil water content continuously changes in response to root water uptake and climatic conditions, potentially resulting in complex system behaviour. In addition, some key variables and parameters are hard (if not impossible) to quantify experimentally. As a result, the whole system is difficult to apprehend, and novel approaches might prove useful to study it.

In the recent years, many computational tools (image analysis, data analysis, process based modelling and upscaling tools) were developed to help quantify and understand water dynamics in the soil-plant system. Some of these tools were developed to work together, or at least be compatible. However, for the uninformed researcher, they might seem disconnected, forming a collection of tools with, at best, a common target (plant-water relation exploration) but unrelated to each other.

The overall objective of this paper is to draw and discuss the role of a functional landscape of interconnected experiments and models for the study of soil-plant water relations. It is articulated as 3 sub-objectives: (i) to inform readers about existing procedures and tools used for the quantification of water flow at the organ, root system and plant population scales, as well as their interconnections forming a comprehensive, though non-exhaustive, network, (ii) to provide examples of studies combining experiments, analytical and modelling tools in this network, motivating the use of such approaches to enhance interpretations of available and future data, and (iii) to identify gaps in the network and argue for a better integration of future tools in this workflow with appropriate experiment and model design. A web interface was developed to help researchers use the network: it is available at https://plantmodelling.shinyapps.io/water_network/.

## Connecting the dots between research tools

Through four examples, we illustrate how the dots, consisting in apparently scattered data and tools, can be connected together in a comprehensive network. These examples span over the different scales mentioned above (organ, root system and population) and for all of them, we present data that can be obtained experimentally, technical limitations that need to be overcome and computational tools readily available. In all these examples, we focus on the soil-root water relation specificities at different scales, except for one where an architectural root growth model is introduced.

### Water flow at the root cross section scale

Different tools and techniques exist to quantify root structural properties at the organ scale. Histology and microscopy techniques enable precise observation of root anatomical structures (the interconnected network of cells). For instance, staining or fluorescence microscopy can be used to acquire images of the organization of different cell types within roots and the nature of cell walls [19]. Different image analysis tools are then available to extract quantitative information out of these images. On the one hand, CellSet [20] is currently the only tool that enables a complete digitization of the entire cell network. As an output, each single cell is represented by a set of connected edges and nodes. Unfortunately, depending on the image quality, the unautomated part of the procedure can be time-consuming. On the other hand, RootScan [21], PHIV-RootCell [22] and RootAnalyzer [23] are fully automated tools that can quantify anatomical properties (such as the number of cells or the mean size of each cell type) but do not provide a digitized cell network.

As a part of the functional layer, cell hydraulic properties are hard to estimate as water fluxes are difficult to measure at this scale. The cell pressure probe enables this estimation from measurements of water pressure relaxation times of individual cells, at a high time cost [4,24]. Osmotic pressure can be measured using nanoliter osmometer [25] or scanning electron microscopy [26]. However, the latter is expensive and generally not part of the standard equipment of a plant physiology laboratory. At the organ scale, the root pressure probe enables the measurement of axial and radial conductivities of root segments [3] and junctions to the stem [27]. Some properties of the system can hardly be determined experimentally such as the partitioning of water pathways across cell layers (apoplastic or cell-to-cell) (Barzana et al, 2012).

Detailed root cross-section anatomical descriptions and a minimal set of empirical cell hydraulic properties (e.g. permeability of cell walls and membranes) enables the simulation of water flow across root cross sections. Like at other scales, water flow in the system is solved using transfer equation with boundary water pressures and conductances as input parameters. Such a model can estimate the equivalent hydraulic conductivity of the root cylinder as well as the partitioning of water flow between apoplastic and symplastic compartments of the system. For instance, by combining measurements of cell and root permeability with a hydraulic model, [28] demonstrated that water flow is primarily apoplastic in lupin roots.

A recent study took advantage of these computational tools to estimate the contribution of pearl millet root types to water uptake. Five types were identified based on cross-section anatomy: primary roots, crown roots and 3 types of lateral roots [29]. A cross-section was thoroughly digitized for each type of root using CellSet [20] (fig. 2A). Root axial hydraulic conductances were estimated using the simplified model of Hagen-Poiseuille (fig. 2B), based on measured xylem vessels dimensions. The digitized root anatomical network served as input for a mechanistic model of radial water flow in roots, namely MECHA [30] (fig. 2C). The model was used to estimate the radial conductivity of a typical segment of each root type. In this example, different tools (image analysis and modelling) were combined to estimate radial and axial conductivities, based on easy-to-acquire experimental data (cross section images). While complementary measurements of root hydraulic properties will always remain an asset (e.g. in order to cross-validate the estimated properties), this method opens the way to high-throughput estimations of root hydraulic properties.

**Figure 2:**
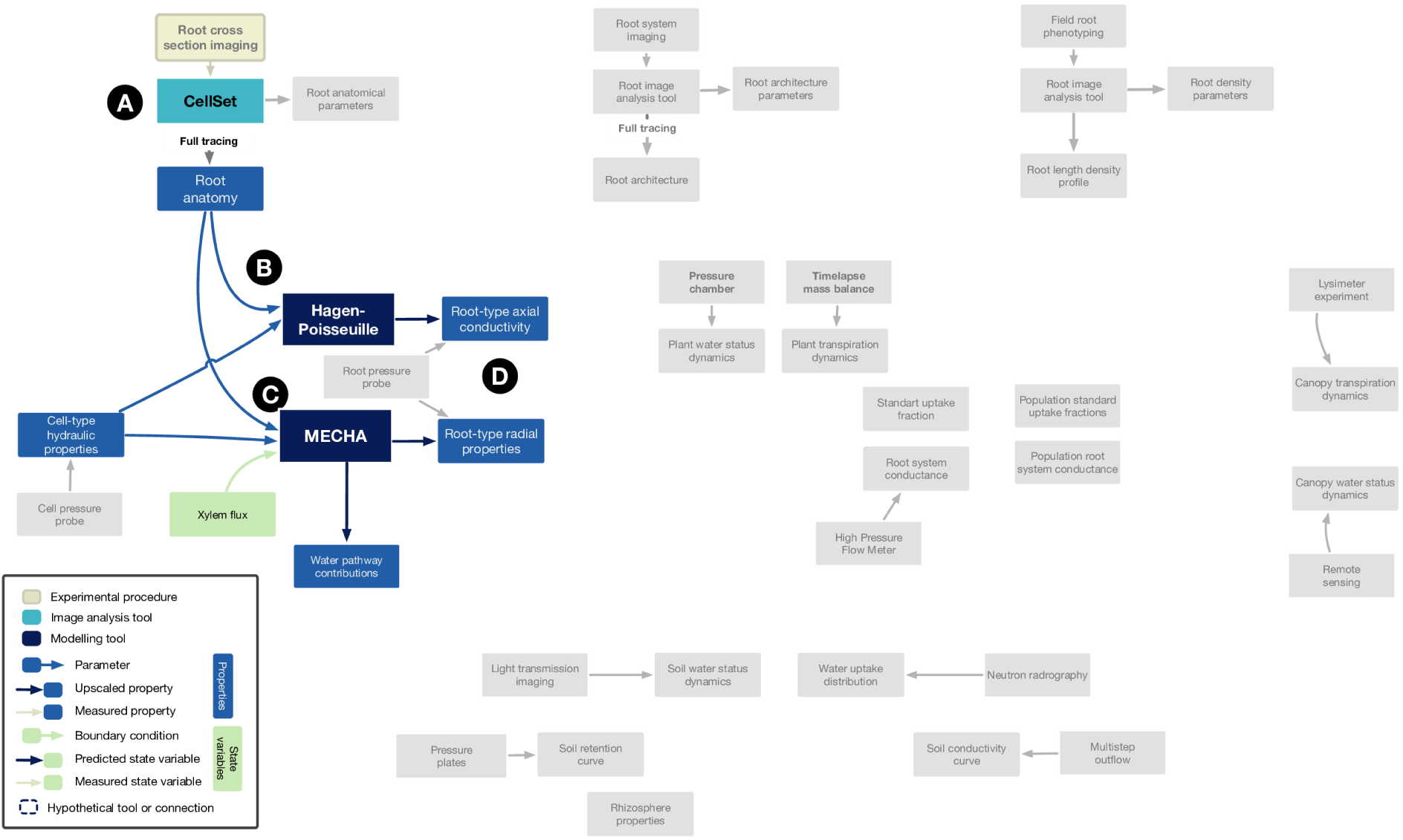
Details of the connected dots to compute the hydraulic properties of the different root types of pearl millet. Colored parts are the tools, models, properties and state variables used in the approach. Specific tools names were added where relevant. See text for details.A: CellSet, B: Hagen-Poiseuille, C: MECHA, D: Output of the different models

### Root system architecture

Unlike plant cells, the root system of annual crops has a convenient macroscopic scale and all elements (roots) are visible to the human eye. However, the main difficulties faced when retrieving the root system architecture are the hidden nature of this part of the plant, the large number of elements that can possibly overlay and the fragility of the smallest roots, making the full excavation of a complete intact root system particularly difficult. Direct manual methods exist to measure single root architectural traits such as the angle of crown roots with a protractor [31] or with the basket method [32,33], the length of individual roots with a ruler [31,34] or a combination of several root architectural traits [31]. However, these manual methods do not give access to the full root architecture.

Several digital tools have been developed and are now widely used to access root architectural traits, mostly from images of root system grown in specific experimental setups (see [35] for a review of existing root phenotyping strategies). These image analysis tools are listed in www.plant-image-analysis.org and will not be detailed here (Lobet et al. 2013, Lobet 2017). The only point to underline is that each tool generally corresponds to a specific growth medium and image capture technique (eg: RooTrak applies to root system growing in 3D and imaged with X-ray Micro Computed Tomography [36]. While some of these tools have been designed to retrace a full root system architecture (often with an important manual input), many of them only extract some root architectural traits (eg. mean lateral root length, number of seminal roots, crown root emergence angle…). Furthermore, even with the use of specifically designed image analysis tools, whole root system digitization becomes time consuming as soon as the plants are several weeks old. Therefore, subsequent tools are needed to reconstruct full root system architectures from extracted root traits.

Root architecture models, such as SimRoot [37], RootBox [38], RootTyp [39], ArchiSimple [40], OpenSimRoot [41] or CRootBox [42], are designed to simulate root systems from a limited number of traits, given as input parameters. The major interest of root architectural models is to generate a large number of contrasted root system architectures. Root system modeling enables the exploration of several variants for the same mean traits and the simulation of contrasted architectures, even from synthetic datasets. These contrasting architectures can then be tested in different scenarios, to identify traits that would be beneficial in challenging environments.

An example that illustrates how root architecture models can be applied to interpret experimental data of other root zone processes is given by Schnepf et al. (2016). Those authors developed a 3D model of the development of mycorrhizal root systems. The model was designed to simulate primary and secondary root infection with arbuscular mycorrhizal (AM) fungi as well as growth of external fungal hyphae in soil. It was calibrated using root architectural data obtained from pot experiments of *Medicago truncatula*, with and without mycorrhizal inoculum of the AM fungus *Rhizophagus irregularis* BEG 158. In those pots, AM root colonization was determined under a compound microscope and the abundance of *R. irregularis* hyphal biomass was determined using real-time PCR. The root system architecture, however, could not be parameterised from those pot experiments. The authors re-used published images from a previous study [43] (fig. 3A) and re-analysed them with the image analysis tool RootSystemAnalyzer [44] (fig. 3B). The traits extracted with RootSystemAnalyzer served as parameters for the RootBox model (fig. 3C) [38] which was used to simulate the root system development of mature plants, together with the arbuscular mycorrhizal fungi. This example highlights how published data can be reused to obtain input parameters for modelling. The current literature is filled with similar resources, opening up numerous opportunities. It also highlights the importance of sharing raw experimental data (in this case the images).

**Figure 3:**
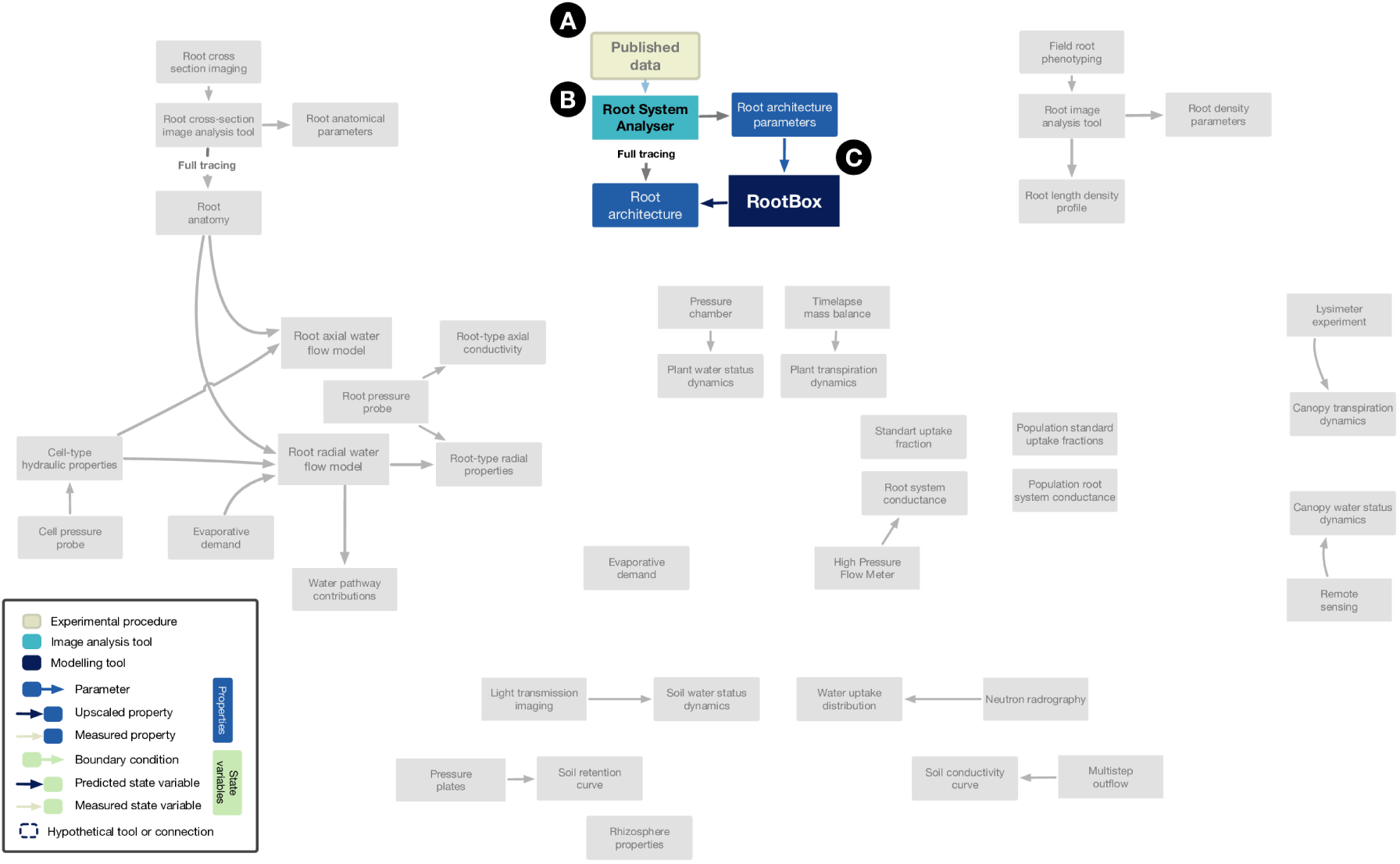
Details of the connected dots for root system architecture generation. Colored parts are the tools, models and properties used in the approach. Specific tools names were added where relevant. See text for details. A: Published data, B: Root System Analyzer, C: RootBox

### Water flow in the root system

At the root system scale, understanding which root traits positively influence plant water uptake dynamics for a given pedo-climatic situation remains an important research question. Ideotypes have been proposed, but are always tied to a specific environment [45–49]. Different traits, either functional or structural, have been found to maximize the final crop yield depending on the environment [50]. Ultimately, we need more than single traits or final yield to have a better understanding of plant-environment interactions. We need to understand how water flow within the plant is dynamically regulated, both spatially and temporally. Unfortunately, accurately measuring water flow is often the limiting step of the experimental pipeline. Several techniques exist to dynamically measure changes in soil water content, such as X-ray computed tomography [51], Electrical Resistivity Tomography [52], neutron tomography [53–56], light transmission imaging [57] or Magnetic Resonance Imaging [58–60]. These techniques can be deployed for a relatively high number of plants. However, due to water capillary flow within the soil domain, observed changes in soil water content are rarely (if never) a direct indication of the location of root water uptake. Water uptake rate itself can be estimated using more advanced but time-consuming lab techniques that use tracers, such as deuterated water that is monitored using neutron radiography [61,62].

Functional-structural plant models (FSPM) are often used to decipher plant-environment relationships [7]. FSPMs couple a complete representation of the root system architecture (or whole plant or shoot) with functional properties. Their input parameters are both functional and structural. For FSPMs simulating soil-root interactions, hydraulic parameters can be obtained using a root pressure probe [63] or the outputs of organ scale models but, as stated earlier, are generally difficult to acquire. Thus they are frequently adapted from the available literature. Rhizosphere hydraulic properties can also be coupled to FSPM [64] and would constitute a central component of plant water availability [16]. Rhizosphere properties are however difficult to parametrize, and would display complex temporal dynamics [65]. The FSPM structural input consists of an explicit representation of the root architecture (see Root system architecture section and the related previous case study). Together with the root system geometry, hydraulic properties define the root system hydraulic architecture [11] and are critical for water stress determination [66,67].

Water-related FSPMs provide a exhaustive description of the root water relations (uptake rates, water potentials, etc.) in both space and time. Thus, they constitute an important way to integrate different types of information about properties of the root system and soil state variables in the root zone, which can be obtained experimentally, and to translate this information into a distributed pattern of water flows and local state variables (e.g. water potentials at the soil-root interface) within the root zone. The latter type of information is, as of today, hardly accessible experimentally. An exhaustive review of FSPMs related to water flow can be found in Ndour et al. [68].

FSPMs can also be used in so-called *inverse modelling* studies. In such case, the output of the model is known and the model is used to estimate one of the input parameters. For instance, the most likely distribution of root hydraulic properties (that are usually assumed to be age- and order-dependent) can be estimated using a soil-root water flow model and laboratory measurements [69]. In this study, measurements of local water fluxes were obtained from neutron radiography at different locations in the root system [56]. As the experiment took place in a rhizotron, the root system could be fully digitized using an appropriate image analysis tool, which provided accurate information on the root system topology and positions in space (fig. 4A and see also Root system architecture section). Water uptake patterns and axial flows within the root system could then be modelled by applying existing water flow equation resolution algorithms [70,71] to the segmented root system (fig. 4B). The water flow model requires boundary conditions that need to be estimated or measured. In this case, the water potential at the root collar was measured using a pressure probe and root-soil interface water potentials were estimated from water content distribution.

**Figure 4:**
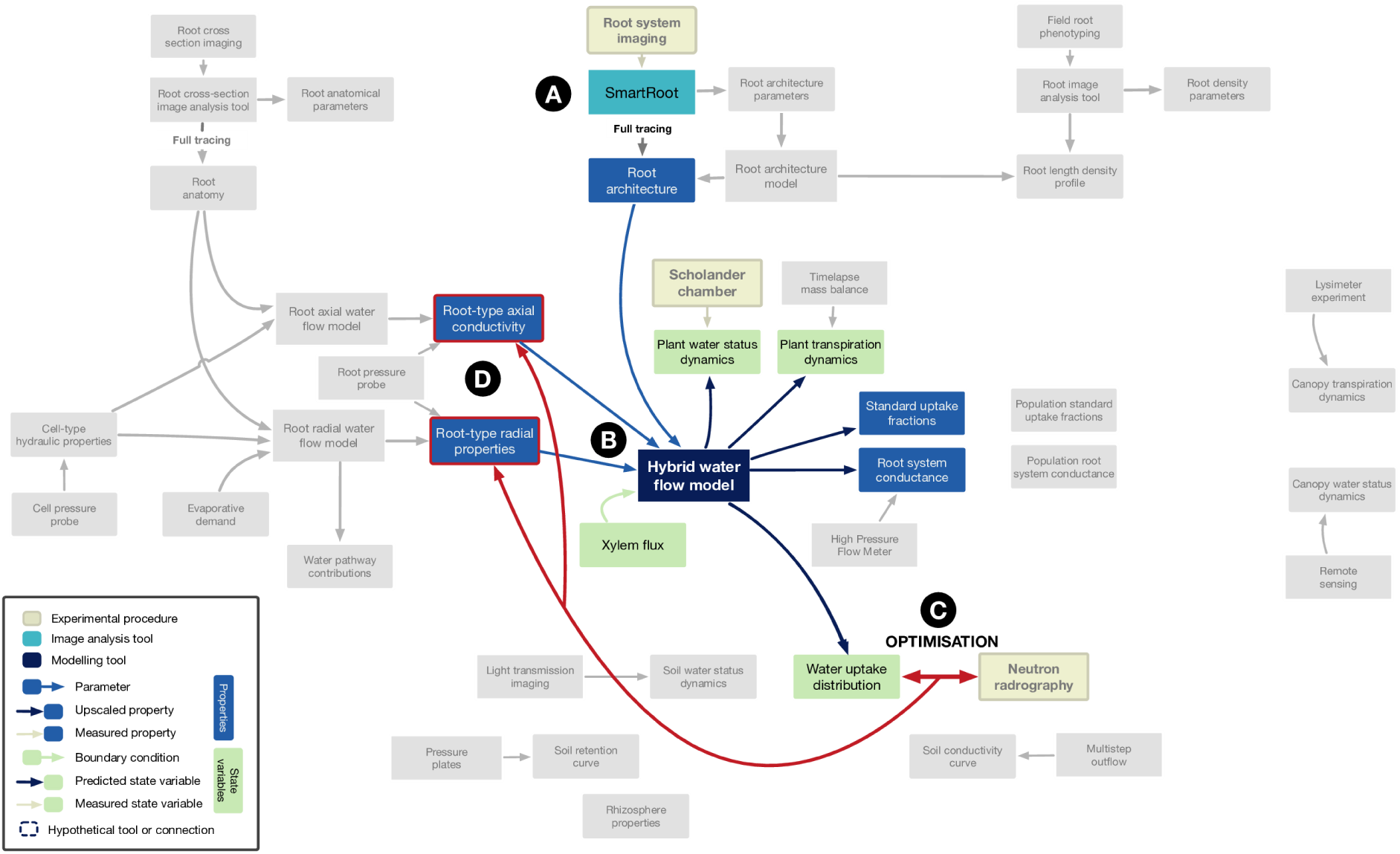
Details of the connected dots for estimating root conductivities from experimental observations through inverse modelling. Colored parts are the tools, models and inputs used in the approach. Specific tools names were added where relevant. The red arrow indicate the optimisation step used in the inverse modelling. The red boxes highlight the variables evaluated using the inverse modelling. See text for details. A: SmartRoot, B: Hybrid water flow model, C: Optimisation, D: Root hydraulic properties

Such a coupling allowed the authors to estimate the parameters of the root hydraulic conductivity function that best fitted the water flow measurements (fig. 4C and D). These parameters then allowed for novel predictions including water uptake and axial flow distributions everywhere in the root system and not only at observed segments, in homogeneous and heterogeneous soil conditions or under various evaporative demands.

### Water flow at the population scale

The population level is a pivotal scale. It interfaces with general circulation models that represent, among others, the circulation of the atmosphere and its interaction with land surface for climate forecasting [72]. It also introduces variables of critical agronomic interest like crop yield per acre [73]. Like at other scales, robust predictions of the system behaviour require the ability to quantify system properties and a proper validation, here involving field scale observations of water fluxes. These fluxes can be estimated with heavy instrumentation and data analytics, for instance using eddy covariance flux towers [74] or soil moisture sensors grids [75].

Structural root information can be obtained using either destructive sampling, such as core sampling [76], monolith excavation [77], trenches [78] or root crown excavation [79], or non-destructive ones, such as minirhizotrons [80]. None of these techniques allows for direct reconstruction of the root system, but rather extract synthetic metrics such as a root length density profile, or root crown data (angles, numbers etc.). Some root architectural traits can be derived from data obtained with these techniques using root architecture models and inverse modeling, as stated above [81,82]. Functional plant properties, such as root system and stomatal conductances, which coordinate shoot water supply [13] and underground water uptake distribution [83], can be characterized on individual plants with low-throughput instruments, such as the high-pressure flowmeter [84] or the porometer [85], then scaled to the population level using the planting density. The main limitation of plant measurement in the field is often the limited sample size, that might not reflect the general behaviour of the system. The same critique can be made about soil hydraulic properties estimated on small and (un)disturbed samples as they may not be representative of the hydrological behaviour at the population level [86,87].

These limitations motivate the use of effective descriptions of population water relations, tailored for this specific scale, such as the transpiration correction for “soil water stress” [88] or one-dimensional soil water and nutrient transfer principles [89]. Two major methodologies address the parametrization of effective field water relations. First, the artificial neural network approach takes advantage of the availability of large amounts of data to train a model. It was used to predict canopy water fluxes from state variables such as the vapor pressure deficit and soil moisture [90,91]. Second, the inverse modelling approach (as described previously) builds on state-of-the-art models to simulate spatio-temporal series of the system state. The model parameter values producing simulated series that best match field observations are considered optimal and representative of the system behaviour. This approach was used to connect models of soil and plant water flow to observations of soil moisture and transpiration in an almond orchard, in order to estimate soil and plant properties, as well as the hardly measurable leaching of water below the root zone [92]. Numerous variables can be used for inverse modeling, such as soil water content, isotopes distributions or root length density profiles.

Going one step further, simplistic macroscale models can be derived from equations of water flow at a lower scale, offering an interesting trade-off between functional simplicity and realism. This type of model involves scale-consistent properties and processes. A cross-validation is thus possible between parameter values estimated directly at the macroscale of interest (e.g. plant population hydraulic conductance per surface area) and derived from the lower scale (e.g. upscaled values derived from root architectural and hydraulic properties). In order to parametrize such a macroscale model of water dynamics in the soil-wheat system, Cai et al. [15] combined one-dimensional process-based models of water flow (i) in soil (Hydrus-1D, [93]) (fig. 5A), (ii) in roots [94], and (iii) in leaves with an isohydric constraint on transpiration [95] (fig. 5B). Regarding soil properties, soil water retention curves were fitted on simultaneous soil water content and pressure head measurements [96] with the software RETC [97] (fig. 5C). However, the parameters of the soil hydraulic conductivity curve were not experimentally determined (fig. 5D). The vertical distribution of roots in the field (root length density profiles over time) was extracted from *in situ* rhizotube pictures, with the software Rootfly [98] (fig. 5E). Its relative distribution is typically used as proxy for the water uptake distribution in uniformly wet conditions [99,100] (fig. 5F). Plant hydraulic properties could not be observed *in situ* for the wheat population (fig. 5F-G).

**Figure 5:**
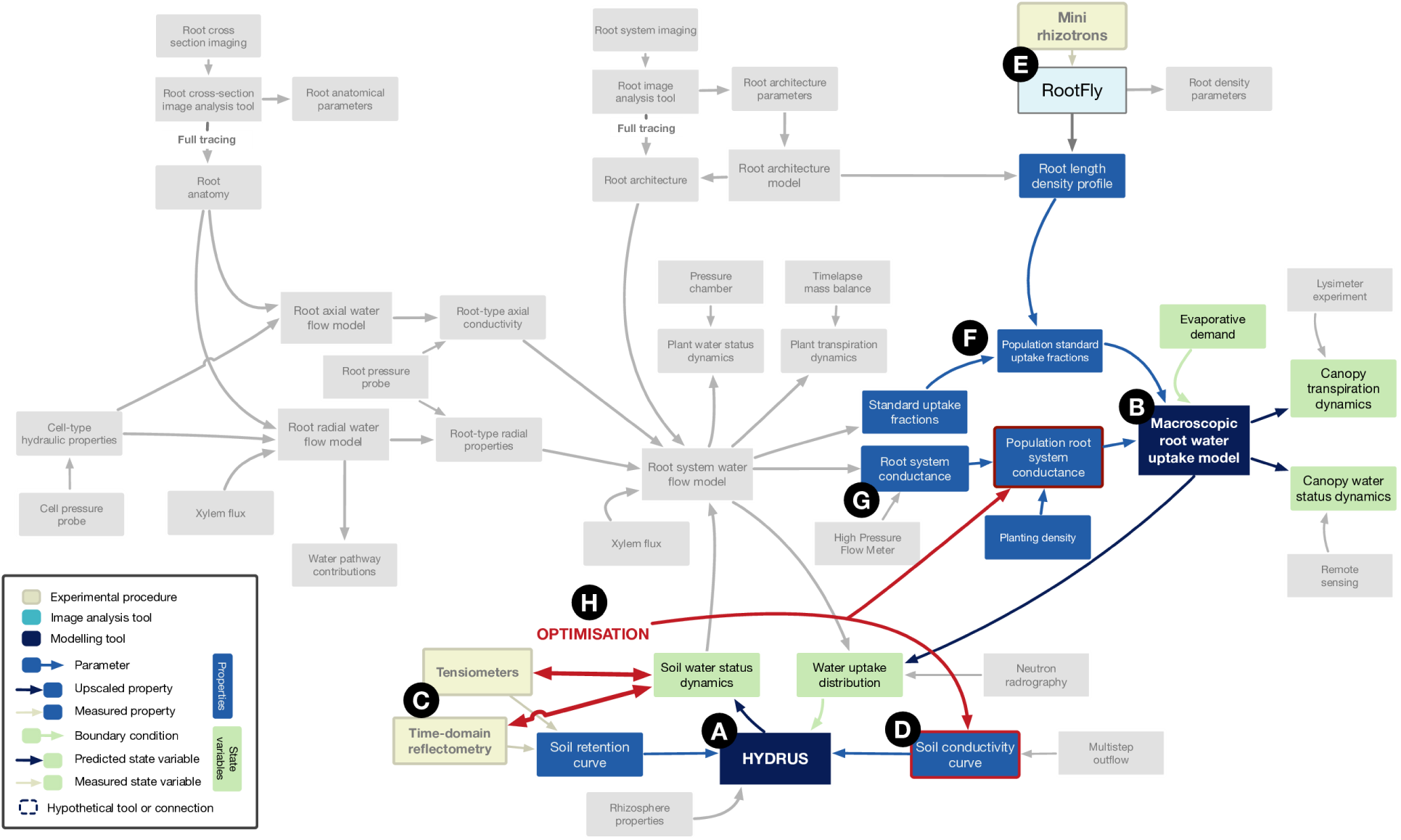
Details of the connected dots for estimated soil and plant-scale conductivities through inverse modelling. Colored parts are the tools, models and inputs used in the approach. Specific tools names were added where relevant. The red arrow indicate the optimisation step used in the inverse modelling. The red boxes highlight the variables evaluated using the inverse modelling. See text for references to letters A-H.

An inverse modelling strategy was therefore used to find the “optimal” soil and plant hydraulic properties (fig. 5H) that best fitted the observed soil water status dynamics. The optimized plant hydraulic parameters were cross-validated with properties at the individual plant scale. Conductance parameters obtained for winter wheat at the same stage of maturity using the hydraulic architecture approach turned out to be consistent with the inversely modeled properties at the population scale [15].

In order to limit the number of parameters, this approach requires the assumption that system properties are time invariant (e.g. soil hydraulic conductivity curve). Because root system conductance tends to scale with root length, the root conductance per unit root length was assumed invariant in order to accommodate for root growth. Such a constraint also matters when accounting for the spatial heterogeneity of root development under different soil/microclimate environments in macroscale simulations [101].

## Discussions and perspectives

Many computational tools exist to better understand water dynamics in soil-plant systems. These tools span different scales (organ, plant and population), types (image analysis, data storage, simulation models) and computational languages (Python, Fortran, C++, C#, Java, …). For the average user, this multitude might seem overly complex and hard to understand. Yet, most of the tools could work together and form a continuous network. Using this network, experimental data can be transferred from scale to scale and generate new insights (fig. 6, and case studies developed above). Modelling tools currently present in the network are listed in table 1. This list is non-exhaustive as the objective of this paper is less to review existing tools than to encourage their integration in order to enhance our understanding soil-plant water relations. For image analysis tools, we refer the reader to the www.plant-image-analysis.org database [102,103].

**Figure 6:**
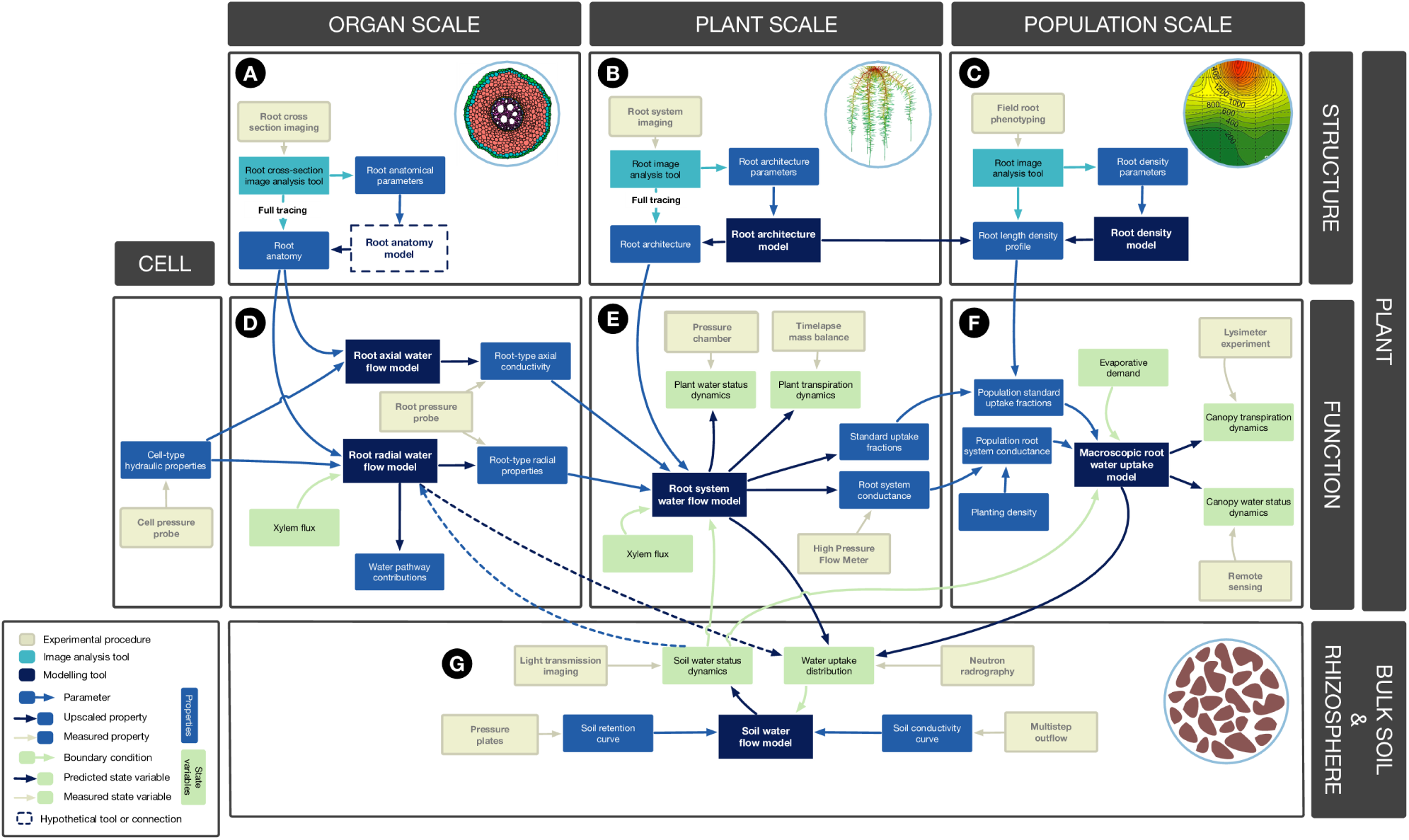
Full network of tools and data used to quantify water flow in the soil plant system. The network connects experimental procedures, computational tools and data related to water flow in the soil-plant system. It is organised by scales (organ, plant and population) and by the types of information (structural or functional, see text for details).A. Tools to quantify the water flow at the organ scale (root section). B. Tools to quantify the structure of root organ. C. Tools to quantify the water flow at the plant scale (root system). D. Tools to quantify the structure of root system. E. Tools to quantify the water flow at the population scale. F. Tools to quantify the structure of root profiles. G. Tools to quantify water flow in the soil.

**Table 1:** List of modelling tools fitting into the network. For image analysis tools, we refer the reader to the www.plant-image-analysis.org website

| Type | Scale | Name | Reference |
| --- | --- | --- | --- |
| Functional | Radial water flow | MECHA | [30] |
| Functional | Root system water flow | R-SWMS | [119] |
| Functional - structural | Root system water flow | PlaNet-Maize | [120] |
| Functional - structural | Root system water flow | OpenSimRoot | [121] |
| Functional | Root system water flow |  | [122] |
| Functional | Soil water flow | HYDRUS | [93] |
| Functional | Soil water flow | RSWMS | [119] |
| Structural | Root system architecture | CRootBox | [123] |
| Structural | Root system architecture | ArchiSimple | [39] |
| Structural | Root system architecture | RootTyp | [124] |
| Structural | Root system architecture | DigR | [125] |
| Structural | Root system density |  | [126] |

We created an interactive online visualization of the network, which contains links to the different tools. We also added a submission form, such as the community could update the network with new (or missing) tools. The online visualisation tool (open-source) is available at https://plantmodelling.shinyapps.io/water_network/

### Identifying gaps in the tool network

Analogies between tool connectivity patterns at each scale in figure 6 reveal the existence of network gaps (represented by the dashed lines in fig.6). Yet, these gaps should not be filled just for the sake of symmetry. Here, we analyze what the function of filling these gaps could be.

The plant “structure” row in Fig. 6 has the most striking pattern, with imaging techniques systematically feeding image-analysis tools. These tools extract two types of data: (i) explicit spatial structures (e.g. RSML, MTG), and (ii) structural pattern properties (e.g. growth rates, branching rates). At the population and plant scales, root development models [39,104] offer the possibility to convert root pattern properties into predicted root structures. While root anatomical patterns can be automatically characterized by image analysis tools such as PHIV-RootCell [22], no root anatomy development model exists at that scale. In the perspective of generating a mechanistic model of a whole plant from the cell scale [105], a root development model would become essential. It would fulfill two main functions: (i) conducting predictions and test hypotheses related to root anatomical development, and (ii) allowing the spatial and temporal interpolation of root anatomies between experimental observations.

Models using explicit root anatomical structures to test hypotheses about hormone signalling [106], tropisms [107] or radial water flow [30] have emerged lately. However, models of axial water flow remain largely underexplored. In the broadly used Poiseuille-Hagen model, only the quantity and diameter of xylem vessels are accounted for. Yet, it is known for a long time that xylem porous plates, pit membranes and persistent primary cross-walls limit root axial conductivity [108–113] and affect the partitioning of water uptake among root types [27].

Experimental methods and numerical tools to represent numerically xylem anatomy and hydraulic properties are missing and may reveal a complexity that is neglected so far. Such numerical representations of xylem vessel structure and hydraulic properties in the axial direction would allow the use of alternatives to Hagen-Poiseuille law [114,115]. Lewis and Boose point out that “Ideally, the exact solutions should be used to calculate [volume flow rate] in xylem conduits, but the equations are difficult to solve without the aid of computer” [115]. Computer availability is no longer an issue and we expect that explicit models of xylem flow will soon emerge. We expect that filling this gap will shed light on the role of cross-walls in the generation of root hydraulic types, and in root-leaf preferential connectivity [116].

Similarly, at the soil-root interface, imaging tools are now available to precisely observe processes at the scale of the soil particle and root hair [117]. Connecting such soil-root interface geometrical descriptions to root hydraulic anatomical models would open new avenues to understand how root hairs enhance plant water availability in dry soils [118].

Soil water fluxes were only explicitly considered in the last case study (population scale). In other case studies, soil was either neglected (organ and plant scales) or included as static boundary conditions. However, in all cases, a model of water flow in the soil domain can be coupled to the plant water flow. Such analyses were for instance carried out at the plant level [41,119,127,128] or the population level [129,130]. Such models may incorporate multiple soil characteristics such as macropores [131], solute convection-dispersion [132] or specific rhizosphere properties [133,134]. For an extensive review of existing soil models, we refer the reader to Vereecken et al. [135].

### Limitations and future developments

Simulating water fluxes in roots with this collection of tools can either help understanding plant water relations as a main goal or be a tool for further application. These tools could also be used as a side usage of a dataset obtained for other purposes. The advent of imaging in plant sciences and the huge progress made in image analysis allow generating high quality quantitative data of plant structure, suitable for model parameterization. Many models exist at different scales and we highlighted the many possibilities to combine these tools. This set of tools greatly increases the potential of interpretation of experimental data. Yet many authors still publish rich datasets without using modelling tools to interpret them. Using models is not trivial for a large part of the plant science community. Coupling several tools and models together is still rarely applied. Several requirements seem essential to facilitate the use of this pipeline whenever it may add value to the data.

A lot of image analysis tools and models flourish in the water transport domain. Thanks to the wide breadth of scientific literature available (scientific papers, reviews, websites…), developers are usually aware of already existing tools and keep in mind to justify the interest of their tool in this landscape. However, we suggest that further efforts should be made to render the tools compatible with existing ones. In this context, the existence of several tools at the same place of the network (eg. RootTyp and CRootBox for root architecture simulation) is not conflicting. Each one can best suit one scale or specific situation. In our opinion, special attention should be paid to the data format. Indeed, the output data format of upstream tools must be compatible with the input format required by downstream ones. If this is not the case, easy-to-handle tools must exist to convert these data. The multiplication of formats and the need to convert data from one type to another may discourage the use of some of the models. In this respect, the existence of standard formats, such as the Root System Markup Language [5] for root architecture, smooths the interconnection between tools.

When a new tool is created, the documentation of its potential connections with existing ones (e.g. in the user guide) would benefit the whole network. It is indeed expected that the knowledge and the use of all modelling tools will increase. It also underlines the need to keep the models and their documentation updated. Pioneering tools sometimes get outdated by new ones that do similar tasks but that are more user-friendly, faster, use the latest formalisms, or are better connected with newly existing tools.

Therefore, either the interconnection between tools needs to be part of a huge maintenance effort for already existing tools, or the acceptance that pioneering tools are doomed to sink into oblivion.

Making the different tools freely available to the community is also a key aspect in their long term maintenance [103]. Many different repositories and licences exist so that everyone should be able to find a combination that suits their (and their institution’s) needs. Free access to the tools’ source codes would indeed greatly facilitate their evolution, reproducibility of the in silico experiments and allow future developers to interconnect them more easily.

## Glossary

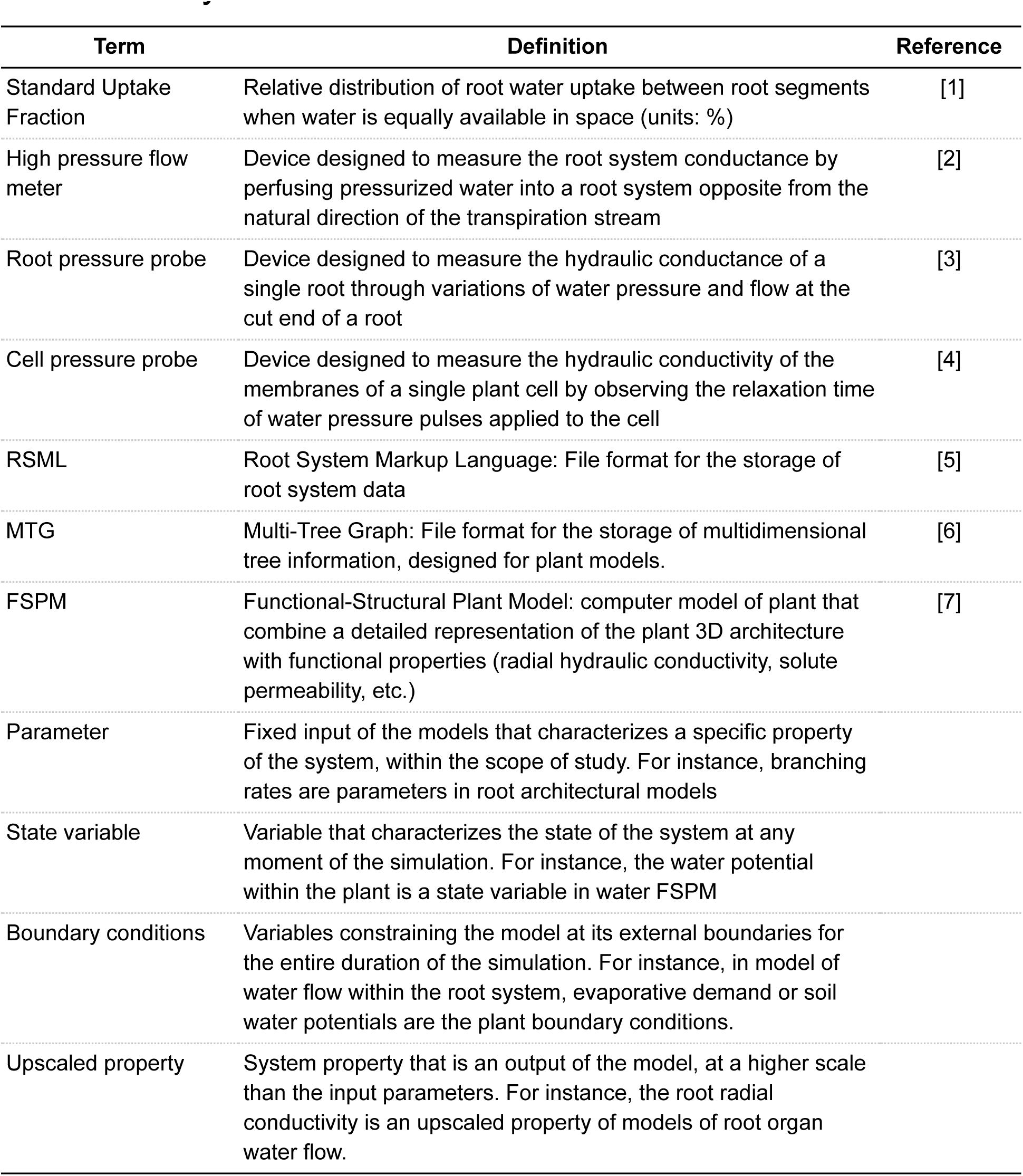

| Term | Definition | Reference |
| --- | --- | --- |
| Standard Uptake Fraction | Relative distribution of root water uptake between root segments when water is equally available in space (units: %) | [1] |
| High pressure flow meter | Device designed to measure the root system conductance by perfusing pressurized water into a root system opposite from the natural direction of the transpiration stream | [2] |
| Root pressure probe | Device designed to measure the hydraulic conductance of a single root through variations of water pressure and flow at the cut end of a root | [3] |
| Cell pressure probe | Device designed to measure the hydraulic conductivity of the membranes of a single plant cell by observing the relaxation time of water pressure pulses applied to the cell | [4] |
| RSML | Root System Markup Language: File format for the storage of root system data | [5] |
| MTG | Multi-Tree Graph: File format for the storage of multidimensional tree information, designed for plant models. | [6] |
| FSPM | Functional-Structural Plant Model: computer model of plant that combine a detailed representation of the plant 3D architecture with functional properties (radial hydraulic conductivity, solute permeability, etc.) | [7] |
| Parameter | Fixed input of the models that characterizes a specific property of the system, within the scope of study. For instance, branching rates are parameters in root architectural models |  |
| State variable | Variable that characterizes the state of the system at any moment of the simulation. For instance, the water potential within the plant is a state variable in water FSPM |  |
| Boundary conditions | Variables constraining the model at its external boundaries for the entire duration of the simulation. For instance, in model of water flow within the root system, evaporative demand or soil water potentials are the plant boundary conditions. |  |
| Upscaled property | System property that is an output of the model, at a higher scale than the input parameters. For instance, the root radial conductivity is an upscaled property of models of root organ water flow. |  |

## Acknowledgments

The authors would like to thank Jennifer Brophy, Heike Lindner and Thérèse LaRue (Carnegie Institution for Science, Stanford, CA) for their useful comment on the first version of manuscript. This work was also supported by the Belgian French community ARC 16/21-075 project. VC was also supported by the Interuniversity Attraction Poles Programme-Belgian Science Policy (grant IAP7/29). During the preparation of this manuscript, FM was supported by the “Fonds National de la Recherche Scientifique” (FNRS) of Belgium as a Research Fellow and is grateful to this organization for its financial support.

## Author contributions

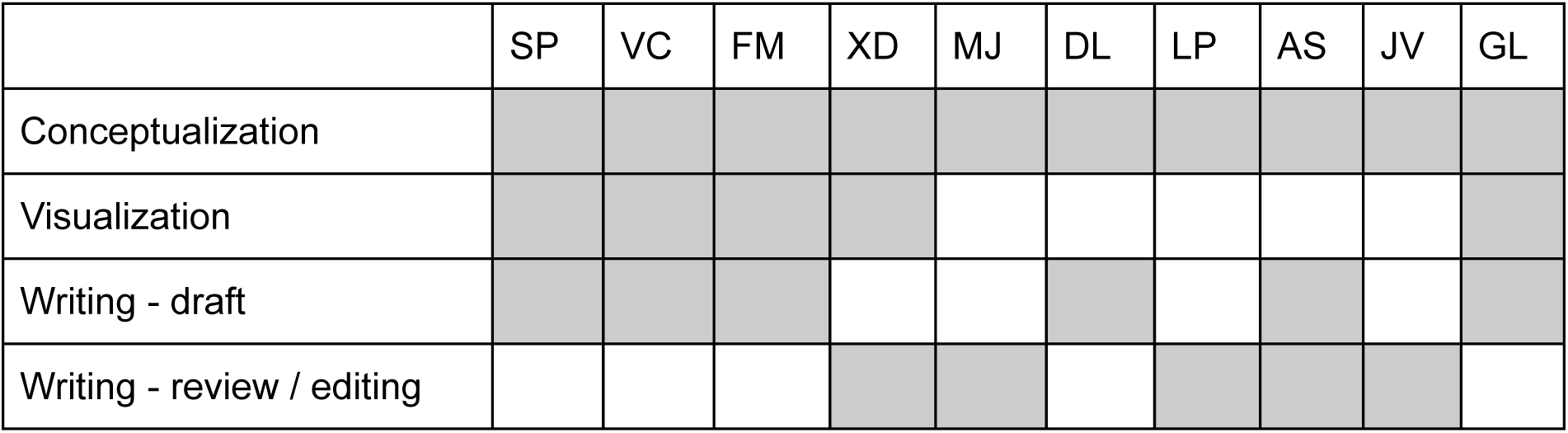

